# *In silico* screening for ERα downmodulators identifies thioridazine as an anti-proliferative agent in primary, 4OH-tamoxifen-resistant and Y537S ERα-expressing breast cancer cells

**DOI:** 10.1101/325746

**Authors:** Claudia Busonero, Stefano Leone, Fabrizio Bianchi, Filippo Acconcia

## Abstract

**Purpose:** Most breast cancers (BCs) express estrogen receptor α (ERα) and are treated with the endocrine therapy (ET) drugs 4OH-tamoxifen (Tam) and fulvestrant (*i.e.*, ICI182,780-ICI). Unfortunately, a high fraction of ET-treated women relapses and become resistant to ET. Therefore, additional anti-BC drugs are needed. Recently, we proposed that the identification of novel anti-BC drugs can be achieved using the modulation of the ERα intracellular content in BC cells as a pharmacological target. Here, we searched for Food and Drug Administration (FDA)-approved drugs that potentially modify the ERα content in BC cells.

**Methods:** We screened *in silico* more than 60,000 compounds to identify FDA-approved drugs with a gene signature similar to that of ICI. We identified mitoxantrone and thioridazine and tested them in primary, Tam-resistant and genome-edited Y537S ERα-expressing BC cells.

**Results:** Mitoxantrone and thioridazine induced ERα downmodulation and prevented MCF-7 cell proliferation. Interestingly, while mitoxantrone was toxic for normal breast cells, thioridazine showed preferential activity toward BC cells. Thioridazine also reduced the ERα content and prevented cell proliferation in primary, Tam-resistant and genome-edited Y537S ERα-expressing BC cells.

**Conclusions:** We suggest that the modulation of the ERα intracellular concentration in BC cells can also be robustly exploited in *in silico* screenings to identify anti-BC drugs and further demonstrate a re-purposing opportunity for thioridazine in primary and metastatic ET-resistant BC treatment.

## 1 Introduction

A shortcut to cancer drug discovery is drug re-purposing, comprising the identification of novel anti-cancer activities of the existing Food and Drug Administration (FDA)-approved drugs. This ‘drug-recycling’ approach reduces the costs for the introduction of novel drugs into the market and bridges the gap between drug discovery and availability [1].

In 70% of all cases, breast cancer (BC) expresses the transcription factor estrogen receptor α (ERα), which is the driver of BC progression. In turn, ERα-positive BC is treated with endocrine therapy (ET) agents that interfere with ERα signaling: the mainstay first-line ET drug is 4OH-tamoxifen (Tam) that binds to ERα and inhibits its transcriptional activity, thus preventing BC progression. However, many Tam-treated women relapse with a resulting metastatic disease resistant to Tam. Tam-resistant BC patients can be treated only with the second-line ET drug fulvestrant (*i.e.*, ICI182,780-ICI), which binds to ERα and induces its degradation, alone or in combination with classic anti-neoplastic drugs [2, 3]. Therefore, novel drugs are needed for the treatment of primary and metastatic Tam-resistant BC.

Analyses of ‘big-data’ derived from omics technologies have facilitated scientific breakthroughs [4, 5]. The iLINCS (Integrative Library of Integrated Network-Based Cellular Signatures) portal is a web platform for the analysis of LINCS signatures. In particular, the iLINCS database contains more than 65,000 gene signatures generated by stimulating diverse cell lines (*e.g.*, BC cells) with different doses of perturbagens (*i.e.*, compounds including FDA-approved drugs) at different time points [6]. Therefore, the iLINCS portal can be used to facilitate the identification of FDA-approved drugs with anti-BC potential.

Recently, we introduced the concept of selective modulation of ERα levels and degradation as an unconventional opportunity to identify anti-BC drugs targeting ERα signaling directly or indirectly. By evaluating the ability of approximately 1,000 FDA-approved drugs to change the ERα intracellular content in BC cells through both classic and high-throughput assays, we identified several compounds with a re-purposing opportunity for primary and metastatic Tam-resistant BC management [7–10].

Here, we used the iLINCS compound database [6] to identify FDA-approved drugs with a predicted activity in modulating ERα and discovered that mitoxantrone and thioridazine induce ERα downmodulation. Moreover, we demonstrate anti-proliferative effects in both Tam-sensitive and Tam-resistant BC cells for thioridazine.

## 2 Materials and Methods

### 2.1 Generation of estrogenic signature in the iLINCS database

To generate an ‘estrogenic signature’ in the iLINCS database, we selected all the signatures available for 17β-estradiol (E2) in MCF-7 cells and all the signatures available for ICI and Tam in MCF-7 cells treated with these drugs for 24 hrs. (Supplementary Table 1). The common 57 genes in the iLINCS signatures upregulated by E2 administration and downregulated by ICI and Tam administration in MCF-7 cells were considered the ‘estrogenic signature’ (Supplementary Table 2; Supplementary Material 1).

### 2.2 Cell culture and reagents

17β-Estradiol (E2), DMEM and fetal bovine serum (FBS) were purchased from Sigma-Aldrich (St. Louis, MO). The Bradford protein assay kit and anti-mouse and anti-rabbit secondary antibodies were obtained from Bio-Rad (Hercules, CA, USA). Antibodies against ERα (F-10 mouse), cathepsin D (H75 rabbit) and pS2 (FL-84 rabbit) were obtained from Santa Cruz Biotechnology (Santa Cruz, CA); anti-vinculin antibody was obtained from Sigma-Aldrich (St. Louis, MO). Chemiluminescence reagent for Western blotting was obtained from Bio-Rad Laboratories (Hercules, CA, USA). Fulvestrant and 4OH-tamoxifen were purchased from Tocris (USA). All the other products were from Sigma-Aldrich. Analytical- or reagent-grade products were used without further purification. The identities of cell lines [*i.e.*, spontaneously immortalized human breast epithelial cells (MCF10a) [11]; human breast carcinoma cells (MCF-7; ZR-75-1; MCF-7 Tam resistant [12]; MCF-7 Y537S ERα mutant [13])] were verified by STR analysis (BMR Genomics, Italy).

### 2.3 Cellular and biochemical assays

Cells were grown in medium supplemented with 10% FBS before any treatment. For E2-based experiments, the cells were grown in 1% charcoal-stripped fetal bovine serum medium for 24 hrs. and then were stimulated with E2 as indicated in the figure captions. For growth curve analysis, cells were plated in 96-well plates in triplicate in growth medium and were counted at the indicated time points after drug administration. Briefly, the medium was removed, and crystal violet solution was added for 20 min at room temperature (R.T.). After extensive washing with tap water, 1% SDS (200 μl) was added to each well to dissolve the crystal violet. The absorbance was read at 595 nm using a Tecan Spark Elisa reader with respect to blank and 0 cells (*i.e.*, crystal violet staining only). The number of cells was derived by comparing the read absorbance with that obtained from a standard curve. Protein extraction was performed as described previously [9]. Western blotting analyses were performed by loading 20-30 μg of protein onto SDS-gels. Gels were run and transferred to nitrocellulose membranes using a Bio-Rad Turbo-Blot semidry transfer apparatus. Immunoblotting was carried out by incubating the membranes with 5% milk (60 min), followed by incubation o.n. with the indicated antibodies. Secondary antibody incubation was continued for an additional 60 min. The bands were detected using a Bio-Rad Chemidoc apparatus.

### 2.4 Cell cycle analysis

After treatment, the cells were grown in medium supplemented with 10% FBS, harvested with trypsin, and counted to obtain 10^6^ cells per condition. The cells were then centrifuged at 1500 rpm for 5 min at 4°C, fixed with 1 ml of ice-cold 70% ethanol and subsequently stained with PI buffer (500 μg/ml of propidium iodide and 320 μg/ml of RNaseA in 0.1% Triton X in PBS). DNA fluorescence was measured using a CytoFlex flow cytometer, and cell cycle analysis was performed using CytExpert v2.0 software (Beckman Coulter).

### 2.5 Affymetrix experiments

Total RNA was extracted using an RNeasy kit (Qiagen) according to manufacturer’s protocol and was quantified using a NanoDrop 2000 system (Thermo Scientific). A GeneChip Pico Reagent Kit (Affymetrix) was used to amplify 5 ng of total RNA according to the manufacturer’s protocol. Quality control of the RNA samples was performed using an Agilent Bioanalyzer 2100 system (Agilent Technologies). Each pool was labeled and hybridized to Affymetrix^®^ Clariom S Human Array according to the manufacturer’s instructions (Affymetrix). The data were normalized using the Signal Space Transformation Robust Multi-array Average (SST-RMA) and Affymetrix Expression Console Software. Gene expression profile analysis was performed using BRB-ArrayTools (v4.6.0) developed by Dr. Richard Simon and the BRB-ArrayTools Development Team. Class comparison analysis to identify differentially regulated genes was performed using t-test with a random variance model and 1000 random data permutations to compute the local false discovery rate.

### 2.6 Statistical analysis

Statistical analysis was performed using ANOVA (one-way analysis of variance and Tukey’s post-test) with the InStat version 3 software system (GraphPad Software Inc., San Diego, CA). Densitometric analyses were performed using the freeware software ImageJ as previously reported [8]. In all analyses, *p* values < 0.01 were considered significant, except densitometric analyses with a chosen threshold of *p* < 0.05.

## 3 Results

### 3.1 *In silico* identification of FDA-approved drugs with an ICI-similar gene signature

Initially, we speculated that drugs interfering with ERα signaling should behave as ICI and Tam in inhibiting the basal levels of those genes induced by E2 administration. Therefore, we generated an ‘estrogenic signature’ in the iLINCS database comprising 57 genes contemporarily upregulated by E2 and downregulated by ICI and Tam (Supplementary Material 1; Supplementary Table 2). Next, we assigned to each gene the variation coefficients and p values obtained by both ICI (Fig. 1a; Supplementary Material 2) and Tam (Fig. 1b; Supplementary Material 3) to produce an ICI- and Tam-specific ‘anti-estrogenic signature’, respectively. These gene signatures were uploaded into the iLINCS database as baits to identify signatures of perturbagen (*i.e.*, drugs) significantly concordant to the ICI- and Tam-specific ‘anti-estrogenic signature’. For both ICI- and Tam-specific ‘anti-estrogenic signatures’, we extracted the perturbagens retrieved for normal breast cells (MCF10a) [11] and ERα-positive (MCF-7) and ERα-negative (MDA-MB-231) BC cells and selected only the MCF-7 cell-specific perturbagen lists (Fig. 1a, 1b, red numbers; Supplementary Table 3 and 4), which, in turn, represent groups of drugs with a gene signature concordant to that of ICI or Tam, only in ERα-positive MCF-7 cells.

**Figure 1.**
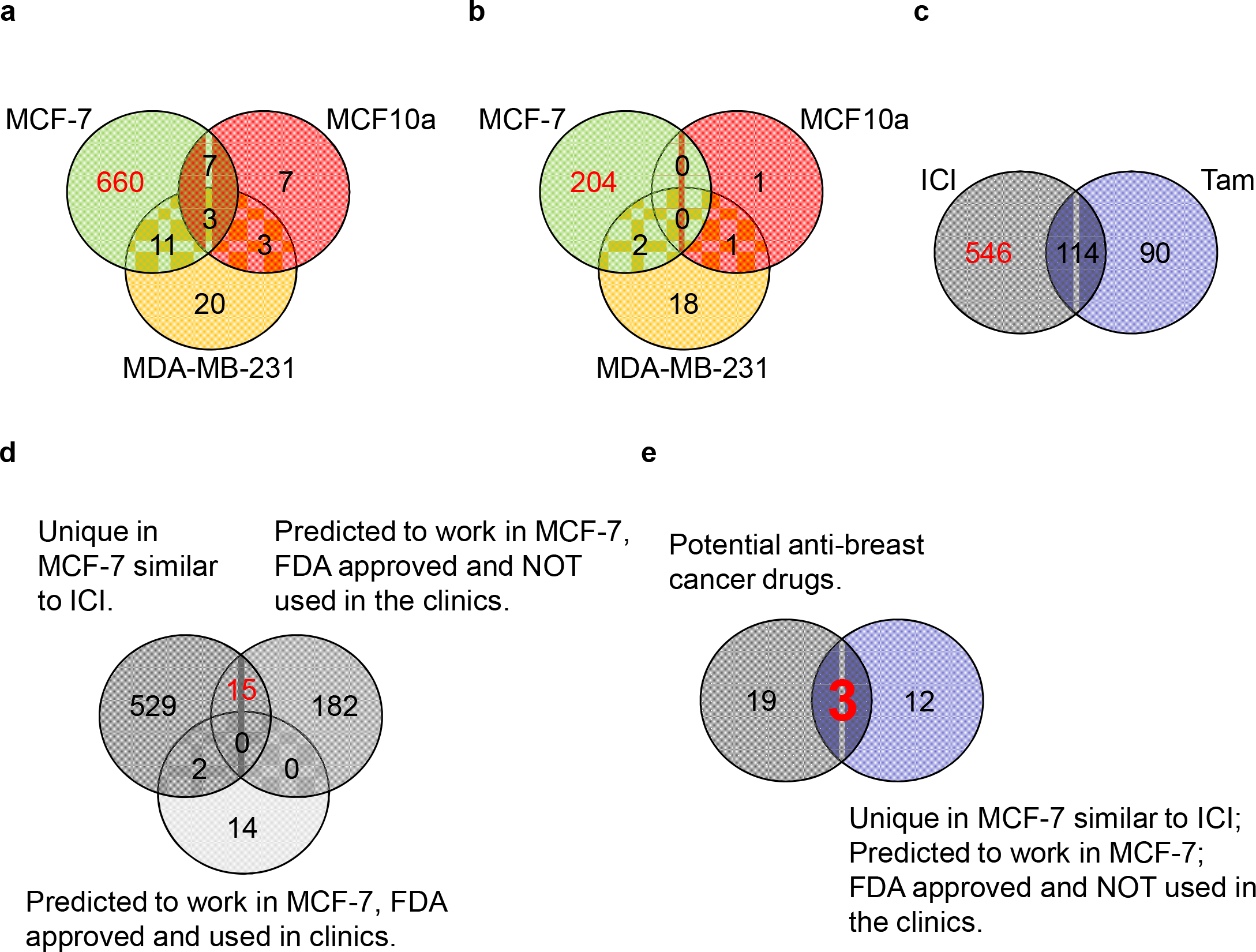
Identification of the candidate FDA-approved drugs not used against BC with ‘antiestrogenic effects’. Venn diagram depicting the number of perturbagens retrieved in the iLINCS database using the ICI (**a**) or Tam (**b**) ‘baits’ in different cell lines. (**c**) Venn diagram depicting the number of perturbagens specific for ICI in MCF-7 cells. (**d**-**e**) Venn diagrams depicting the final choice of FDA-approved drugs not used against BC with ‘anti-estrogenic effects’. Red numbers indicate the perturbagen chosen at each step of the identification procedure. For details, please see the text.

Next, we searched for drugs potentially inducing ERα downmodulation by assuming that such compounds should have generated a gene signature concordant only to that of ICI. Thus, we compared the ICI- and Tam-concordant MCF-7 cell-specific lists of perturbagens and exclusively selected those that were ICI specific (Fig. 1c, red number; Supplementary Table 5). The resulting 546 perturbagens were filtered out by those drugs previously predicted to work as anti-cancer drugs in MCF-7 cells, FDA approved and used or not in the clinics for BC treatment [14] (Fig. 1d). Next, we selected only those 15 drugs with i) a gene signature concordant to that of ICI only in MCF-7 cells, ii) predicted to work as anti-cancer drugs in MCF-7 cells and iii) FDA approved but iv) not used in the clinics for BC treatment (Fig. 1d; Supplementary Table 6). Finally, we intersected our 15 candidate drugs with 22 drugs identified *in silico* as repurposing candidates for BC therapy [15]. Figure 1e indicates mitoxantrone, thioridazine and menadione as common candidate anti-BC compounds.

### 3.2 Validation of mitoxantrone and thioridazine as drugs inducing ERα downmodulation and preventing proliferation in cells modeling primary BC

According to our reasoning, mitoxantrone, thioridazine and menadione should induce ERα degradation and prevent proliferation in MCF-7 cells in a manner similar to ICI. Already published data have demonstrated such activities for menadione in BC cells [16, 17]. Interestingly, we observed that 24 hrs of administration of mitoxantrone and thioridazine to MCF-7 cells induced a reduction in the ERα intracellular content (Fig. 2a, 2a’) and a cell cycle block with a significative reduction of S phase, at the expense of the G2 and G1 phases for mitoxantrone and thioridazine, respectively, while as expected, ICI treatment of MCF-7 cells induced ERα degradation and an accumulation of the cells in the G1 phase of the cell cycle (Fig. 2a, 2a’, 2b, 2b’). To confirm the proliferative block, we also measured the cell number after 3 days of administration of different doses of mitoxantrone, thioridazine and ICI in MCF-7 cells. As control of the potential toxicity of the drugs in non-cancerous cells, parallel experiments were performed in MCF10a cells, which are a model of spontaneously immortalized human breast epithelial cells [11]. Figure 2c indicates that ICI reduced the number of MCF-7 cells at all tested doses, but it did not affect the proliferation of MCF10a cells. Conversely, mitoxantrone administration determined the same dose-dependent linear reduction in both the MCF-7 and MCF10a cell number (Fig. 2d). Interestingly, thioridazine decreased the number of MCF-7 cells in a dose-dependent manner but significantly reduced MCF10a only at the highest dose tested (Fig. 2e). These data indicate that mitoxantrone and thioridazine reduce the ERα intracellular levels and prevent proliferation in MCF-7 cells in a manner similar to ICI as predicted by our *in silico* analysis.

**Figure 2.**
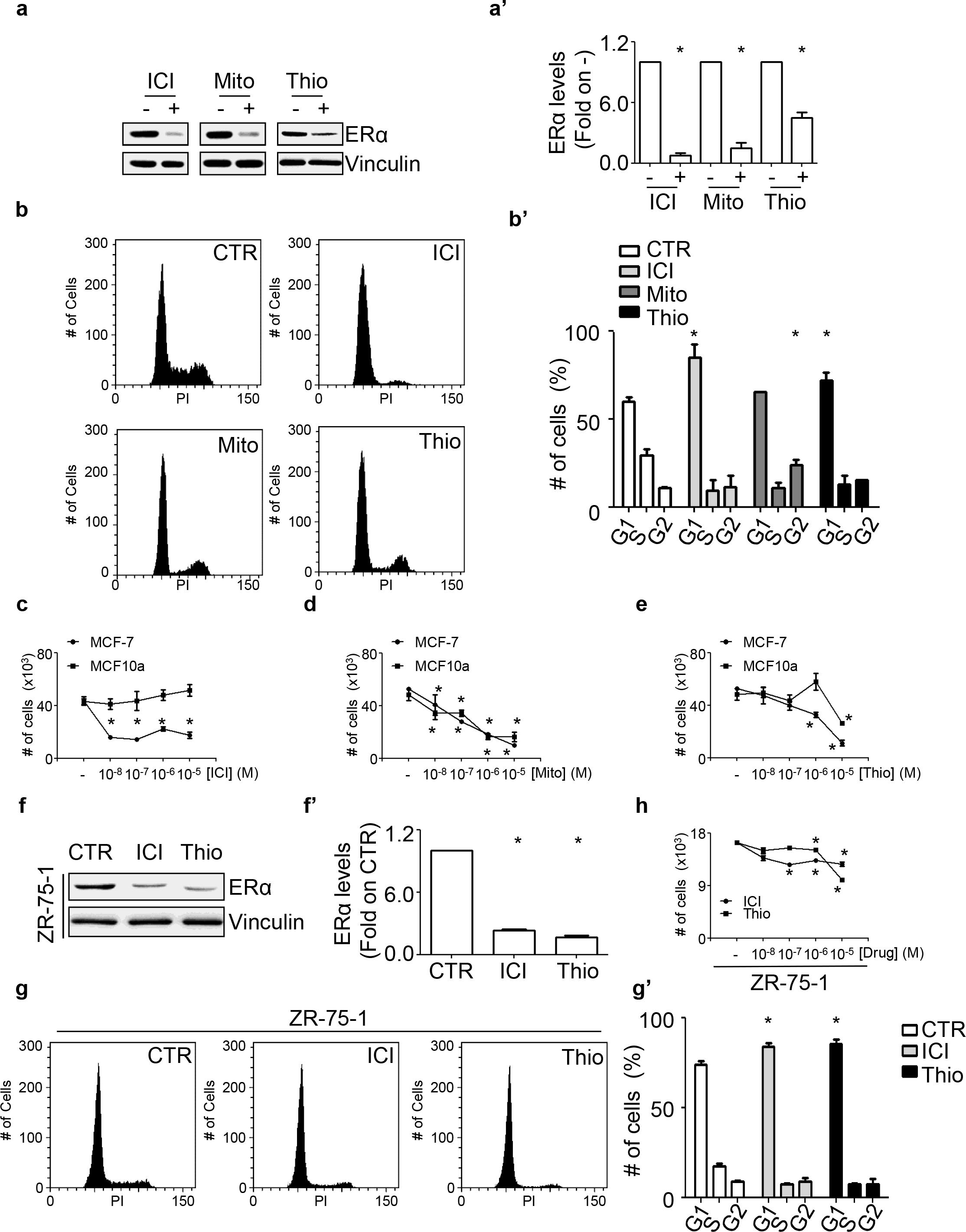
Effect of mitoxantrone and thioridazine on the ERα levels and cell proliferation in MCF-7 and ZR-75-1 cells. Western blotting and relative densitometric analyses of the ERα expression levels in MCF-7 cells **(a, a’)** treated for 24 hrs. with either ICI182,780 (ICI; 0.1 μM), mitoxantrone (Mito; 0.1 μM), thioridazine (Thio; 10 μM) or vehicle (DMSO). The loading control was represented by vinculin expression in the same filter. Cell cycle analyses in MCF-7 **(b, b’)** cells treated as described in **(a)**. Number of MCF-7 and MCF10a cells treated at the indicated doses for 3 days with either ICI182,780 (ICI) **(c)**, mitoxantrone (Mito) **(d)**, or thioridazine (Thio) **(e)**. **(f, f’)** Western blotting and relative densitometric analyses of the ERα expression levels in ZR-75-1 cells treated for 24 hrs. with either ICI182,780 (ICI; 0.1 μM), thioridazine (Thio; 10 μM) or vehicle (DMSO; - CTR). The loading control was represented by vinculin expression in the same filter. Cell cycle analyses in ZR-75-1 cells **(g, g’)** treated as described in **(f)**. **(h)** Number of ZR-75-1 cells treated at the indicated doses for 3 days with either ICI182,780 (ICI) or thioridazine (Thio). * indicates significant differences with respect to the control (DMSO; - CTR) sample.

Because mitoxantrone significantly affected the proliferation of non-cancerous MCF10a cells at all the doses tested, we focused on thioridazine. Next, experiments were performed to demonstrate the effectiveness of thioridazine in other ERα-expressing BC cells. ZR-75-1 cells were treated for 24 hrs with thioridazine and ICI, and both the levels of ERα and the number of cells in the different phases of the cell cycle were measured. Figure 2f, 2f’, 2g and 2g’ show that both thioridazine and ICI induced a 90% reduction in the ERα content and increased the number of cells in the G1 phase of the cell cycle. In particular, each drug increased the G1 content from 74% to 85%. Accordingly, thioridazine also reduced the number of ZR-75-1 cells in a dose-dependent manner (Fig. 2h). These data indicate that thioridazine demonstrates anti-proliferative activity towards cell lines modeling primary BC and open the possibility that this drug could be effective also in cell lines modeling metastatic BC cells.

### 3.3 Analysis of thioridazine as a drug inducing ERα downmodulation and preventing cell proliferation in cell lines modeling metastatic BC

4OH-Tamoxifen resistance is a critical issue in BC management. Thus, novel drugs inducing ERα degradation in Tam-resistant (Tam-Res) metastatic BC are demandingly required [18]. Because we found that thioridazine reduces the ERα intracellular content and preferentially affects cancer cell proliferation over normal breast cells (Fig. 2), we next analyzed the impact of thioridazine on ERα levels and proliferation of both Tam-Res MCF-7 cells, which were previously selected by continuous administration of 100 nM Tam [12], and genome-edited Y537S ERα MCF-7 (Y537S-MCF-7) cells, which express a point-mutated form of ERα that is most frequently found in relapsing metastatic BC. The expression of this receptor variant confers to tumor cells resistance to Tam, E2 insensitivity and ERα resistance to ICI-induced degradation [13, 19–21].

Initial experiments were performed to characterize these cellular models of metastatic BC. The growth of Tam-Res MCF-7 and Y537S-MCF-7 cells was analyzed under different doses of Tam treatment for 7 days compared with that of both MCF-7 and ZR-75-1 cells. As shown in Supplementary figure 1a, Tam induced a dose-dependent reduction in the cell numbers of both the MCF-7 and ZR-75-1 cell compared with that of Tam-Res MCF-7 and Y537S-MCF-7 cells, which were not affected by Tam at any of the doses tested. Next, we performed Affymetrix analysis to compare the gene expression profile of the Y537S-MCF-7 cell line with that of MCF-7 cells.

Differentially expressed genes that were significantly upregulated (FDR<5%) in Y537S-MCF-7 compared with that in MCF-7 cells were further compared with those upregulated genes identified in two other published datasets [13, 19,20]. Among the three lists, 19 genes were commonly found to be upregulated (Supplementary figure 1b and Supplementary Table 7). Interestingly, 17 of the 19 genes were also upregulated by E2 in MCF-7 cells [13] and were previously known E2-target genes (*e.g.*, presenilin2, pS2; cathepsin D, CatD; caveolin-2) [22]. Accordingly, the basal pS2 and CatD levels were strongly upregulated in Y537S-MCF-7 cells, and E2 was less effective in increasing the pS2 and CatD intracellular levels in MCF-7-Y537S than in MCF-7 cells (Supplementary Fig. 1c). Furthermore, Y537S-MCF-7 cells lost the ability to proliferate in response to E2 administration (Supplementary Fig. 1d). Therefore, notwithstanding the difference in the culturing conditions [13, 19,20], these data confirm that the ERα Y537S mutation confers Tam resistance and E2 insensitivity to cells and strongly support Tam-Res MCF-7 and Y537S-MCF-7 cells as a valuable model mimicking metastatic BC [13].

Next, we measured the effect of thioridazine on the ERα levels in both Tam-Res MCF-7 and Y537S-MCF-7 cells. ICI was also included in the experiment as a control for ERα degradation. Figure 3a, 3a’ 3d and 3d’ show that 24 hrs of thioridazine administration induced a reduction in the ERα intracellular content by more than 90% in both cell lines. Notably, the thioridazine effect on ERα levels was more pronounced than that obtained by ICI administration (Figure 3a, 3a’ 3d and 3d’). We next evaluated the impact of thioridazine and ICI on cell cycle perturbation in both Tam-resistant cell lines. Figure 3c, 3c’, 2g and 2g’ show that both thioridazine and ICI increased the number of cells in the G1 phase of the cell cycle with thioridazine, leading to an increase in the G1 content up to 77% in both cell lines. These data indicate that thioridazine reduces the ERα cellular content and blocks the cell cycle in the G1 phase in both Tam-Res MCF-7 cells and Y537S-MCF-7 cells.

Next, to confirm the proliferative block, we counted the number of both Tam-Res MCF-7 and Y537S-MCF-7 cells after 3 days of administration of different doses of thioridazine and ICI. Figure 3b shows that, in Tam-Res MCF-7 cells, both ICI and thioridazine induced a dose-dependent reduction in the cell number. By contrast, ICI (at doses higher than 10^−7^ M), but not thioridazine, reduced the number of Y537S-MCF-7 cells (Fig. 3e).

**Figure 3.**
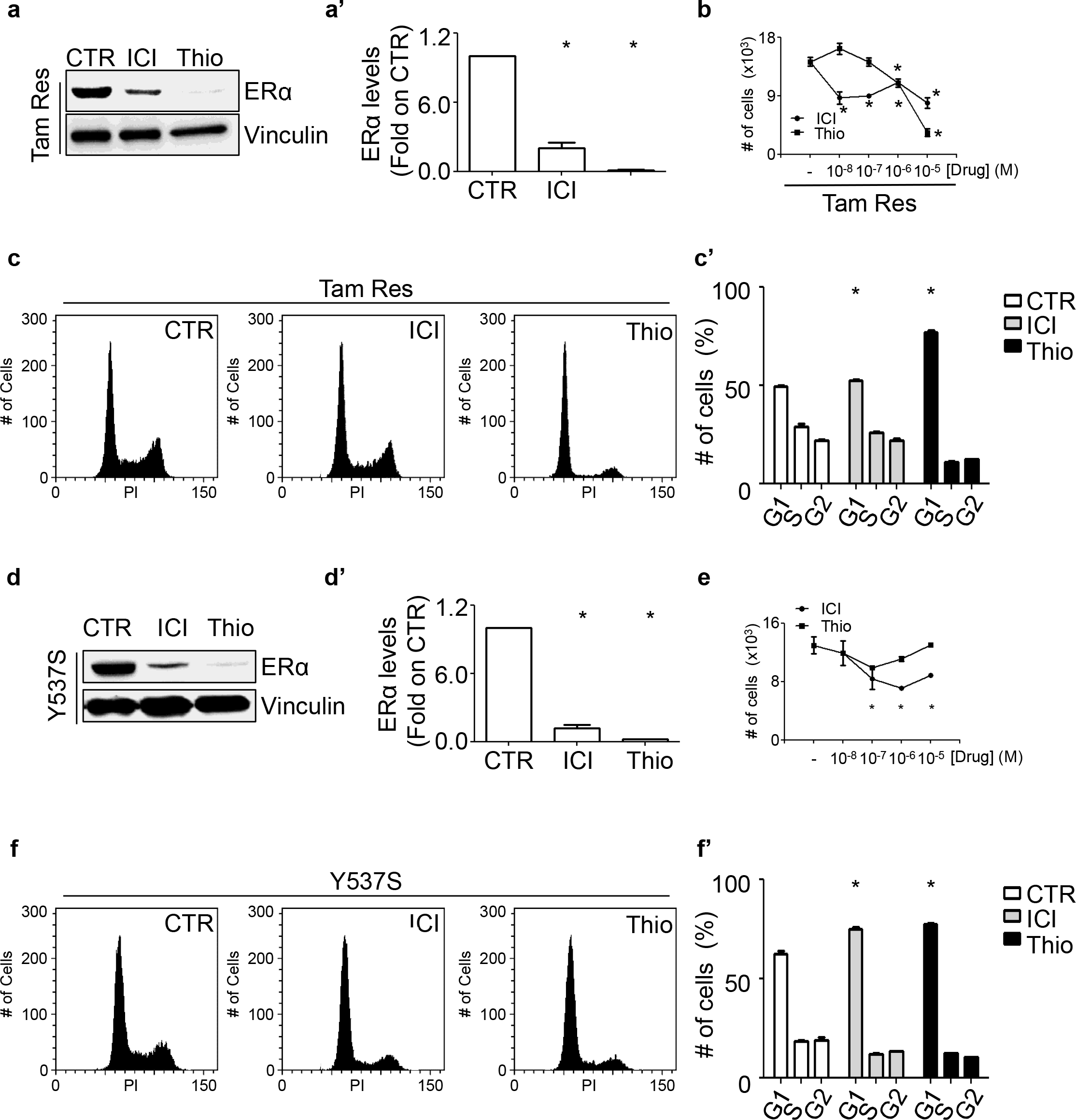
Effect of thioridazine on the ERα levels and cell proliferation in Tam-Res MCF-7 and Y537S ERα-expressing MCF-7 cells. Western blotting and relative densitometric analyses of the ERα expression levels in Tam-Res **(a, a’)** and Y537S ERα-expressing MCF-7 cells **(d, d’)** treated for 24 hrs. with either thioridazine (Thio; 10 μM), ICI182,780 (ICI; 0.1 μM) or vehicle (DMSO; - CTR). The loading control was represented by vinculin expression in the same filter. Number of Tam-Res **(b)** and of Y537S ERα-expressing MCF-7 cells **(e)** treated at the indicated doses for 3 days with either ICI182,780 (ICI) or thioridazine (Thio). Cell cycle analyses in Tam-Res **(c, c’)** and Y537S ERα-expressing MCF-7 cells **(f, f’)** treated as described in **(a, d)**. * indicates significant differences with respect to the control (DMSO; - CTR) sample.

### 3.4 Analysis of ICI and thioridazine combination treatment in BC cells

Metastatic BC can be only treated with limited pharmacological options (*e.g.*, ICI) [23]. Consequently, different lines of research are trying to identify novel molecules that induce ERα degradation in Tam-resistant metastatic BC [18, 24,25] or new pathways that could be targeted alone or in combination with ICI [13, 19, 20, 26, 27]. Therefore, we next asked whether the combination of ICI and thioridazine could exhibit anti-proliferative synergism in primary and metastatic BC cell lines. In turn, we treated MCF10a, MCF-7, ZR-75-1 and Tam Res-MCF-7 cells with ICI and thioridazine alone or in combination for 3 days (Fig. 4a). The MCF10a cell number did not change with any of the indicated treatments. By contrast, the numbers of MCF-7 cells, ZR-75-1 cells and Tam-Res MCF-7 cells were significantly reduced upon ICI and thioridazine treatment (Fig. 4a). Interestingly, ICI and thioridazine co-treatment had a slightly but significantly stronger effect than the drugs alone only in MCF-7 cells and in Tam-Res MCF-7 cells (Fig. 4a).

**Figure 4.**
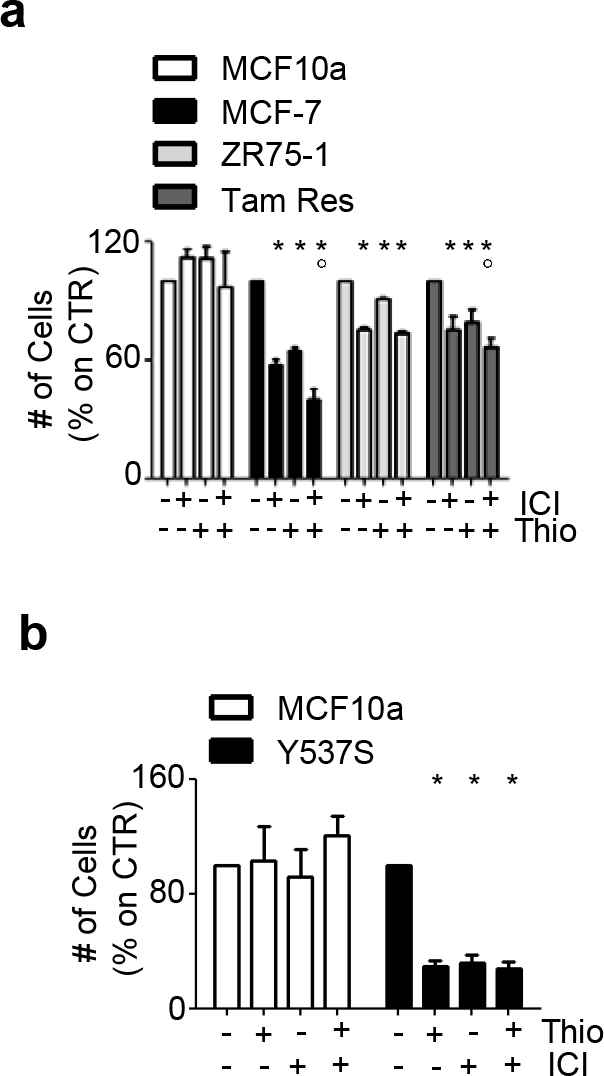
Effect of thioridazine and ICI182,780 co-treatment in BC cells. **(a)** Number of MCF10a, MCF-7, ZR-75-1, and Tam-Res MCF-7 cells treated for 3 days with either ICI182,780 (ICI; 0.1 μM) and thioridazine (Thio; 1 μM) alone or in combination. **(b)** Number of Y537S ERα-expressing MCF-7 (Y537S) and MCF10a cells treated for 7 days with either ICI182,780 (ICI; 0.1 μM) and thioridazine (Thio; 1 μM) alone or in combination. * indicates significant differences with respect to the control (DMSO; - CTR) sample. ° indicates significant differences with respect to the control ICI and Thio samples.

Next, we repeated the same experiments in Y537S-MCF-7 cells and found that the effect of ICI on cell proliferation was reduced (Fig. 3e) compared with that in MCF-7 cells (Fig. 2c). Therefore, we reasoned that such reduced drug sensitivity could occur also for thioridazine, and, indeed, when the Y537S-MCF-7 cells were treated with ICI and thioridazine alone or in combination for 7 days, both ICI and thioridazine reduced the cell number to the same extent, while their co-administration did not further decrease the Y537S-MCF-7 cell number (Fig. 4b).

Altogether, these data indicate that thioridazine has an anti-proliferative activity in cell lines modeling both primary and metastatic BC.

## Discussion

The main aim of the present work was to challenge the concept that the modulation of the ERα intracellular concentration is a signaling parameter that can be used to identify anti-BC drugs. We developed this notion because of the following: i) Tam stabilizes ERα, ICI destabilizes it, and both receptor ligands inhibit BC cell proliferation; and ii) molecules not necessarily binding to ERα or genetic interferences with cellular pathways unrelated to ERα can increase/decrease the ERα content in BC cells and prevent cell proliferation [10]. Therefore, compounds altering the ERα protein amount in BC cells can, in principle, prevent BC cell proliferation. Consequently, we devised a strategy in which, by applying a library of approximately 1,000 FDA-approved drugs to BC cells, we could identify molecules possessing as a new activity (*i.e.*, ‘drug re-purposing’ approach) the capability to modify ERα protein levels and inhibit cell proliferation. By using different assays, we discovered that several FDA-approved drugs (*e.g.*, emetine, carfilzomib and methotrexate) change the ERα protein levels and prevent BC cell proliferation through different mechanisms [7–9].

Here, we exploited a library of more than 60,000 compounds annotated in the iLINCS database [6] to identify *in silico* FDA-approved drugs that could modify the ERα content and prevent cell proliferation in BC cells. Thus, we developed an ‘anti-estrogenic signature’, which we used as a bait to fish out similar genetic signatures to Tam and ICI within the entire iLINCS dataset only in ERα-expressing MCF-7 cells (Fig 1a and 1b). Although Tam is the mainstay of clinical treatment for ERα-positive breast cancers [23], approximately 40% of all ERα-positive BC cases treated with Tam relapse as metastatic disease, which is then treated with fulvestrant (*i.e.*, ICI) to induce ERα degradation. Because fulvestrant has a poor oral activity and possesses diverse side effects [23], we next searched for a list of perturbagens specific for MCF-7 cells with a predicted activity similar to that of ICI, and then we intersected these with other *in silico* analyses [14, 15] of FDA-approved drugs with anti-BC potential. In this way, we identified mitoxantrone, thioridazine and menadione, which indeed were shown to reduce the ERα intracellular content in both MCF-7 and ZR-75-1 cells.

Validation of the mitoxantrone and thioridazine effect was next performed in two ERα-expressing BC cells (*i.e.*, MCF-7 and ZR-75-1) mimicking primary BC compared with a widely-used model of breast cells mimicking normal breast epithelial cells [11]. In both BC cell lines, either drug reduced the ERα intracellular content. Control of the ERα intracellular concentration in BC cells is critical for their growth, survival and spreading. Therefore, the ERα abundance is regulated by different mechanisms acting at both the transcriptional and post-transcriptional levels [10]. Although we did not aim to identify the underlying mechanisms for the ability of mitoxantrone, a topoisomerase II inhibitor that induces DNA damage [28], and of thioridazine, a drug used to treat psychiatric disorders [29], to reduce ERα protein levels, our *in silico* analysis revealed additional activities for these already FDA-approved drugs that deserve future detailed investigations. Nonetheless, based on the present data and because menadione can reduce the ERα intracellular content [16, 17], we conclude that the modulation of the ERα intracellular concentration in BC cells is a robust pharmacological target that can be used also for *in silico* screenings to fish out drugs affecting the ERα content in BC cells.

As noted above, only limited pharmacological options are available for metastatic Tam-resistant BC that still express ERα [23]. There are two main classes of Tam resistance: *de novo* resistance refers to tumors that do not respond to Tam treatment from the beginning, while *acquired* resistance refers to neoplasia that initially respond to Tam and usually, within a period of 15 years, relapse and become insensitive to Tam treatment [30]. Many mechanisms have been reported for the development of Tam resistance in BC [31]. However, recently, it has been shown that a significant fraction of metastatic BC express a mutated, constitutively active variant of ERα (*e.g.*, Y537S ERα), which confers to the disease hormone-independent growth, Tam insensitivity and resistance to ICI-induced ERα degradation [18]. In turn, new compounds that target ERα in metastatic BC as additional ET drugs are strongly required, and, indeed, new orally active antiestrogens that directly bind to and induce the degradation of the mutated ERα form [18, 24,25] are being searched.

Therefore, the above-described results prompted us to test the ability of thioridazine to influence the ERα intracellular content in two cellular models mimicking metastatic BC (*i.e.*, Tam-resistant MCF-7 cells [12] and genome-edited MCF-7 cells expressing the Y537S ERα mutant [13]). Remarkably, we report that thioridazine reduces the ERα content in the cell lines mimicking metastatic BC (Fig 3). In this respect, it is important to stress here that thioridazine reduces the intracellular content of the Y537S ERα mutant, which is resistant to ICI-induced degradation [13, 18], especially considering that this receptor variant completely transforms the landscape of the gene expression pattern in the metastatic cellular context [32]. In turn, the identification of drugs eliminating the driver of such a genetic drift in metastatic disease is paramount. Here, we propose thioridazine as one of such compounds. As pointed above, we did not study the molecular mechanisms by which thioridazine reduces the ERα content; thus, under no circumstances do we imply to refer to thioridazine as a novel selective estrogen receptor downmodulator (SERD). Nonetheless, this FDA-approved drug used to treat psychiatric disorders phenocopies the effects of the prototype SERD fulvestrant and, thus, can be added to the list of drugs inducing ERα degradation in metastatic BC cells.

According to the modification of the ERα intracellular concentration in BC cells [7–10], mitoxantrone, thioridazine (Fig. 2) and menadione [16, 17] interfere with BC cell proliferation. In particular, mitoxantrone and thioridazine block the cell cycle and reduce the cell number in a dose-dependent manner. Ideal anti-cancer drugs should only kill malignant cells. However, mitoxantrone seems toxic for both normal and BC cells (Fig. 2d) as could be expected by a DNA-damaging compound. By contrast, thioridazine preferentially targets primary and metastatic BC cells. Therefore, our data indicate that the reduction in the ERα intracellular levels could be regarded as a possible anti-proliferative mechanism for thioridazine action in BC cells. Moreover, although additional studies are needed to define the toxicity profiles of thioridazine to establish correct therapeutically protocols for BC treatment, the anti-proliferative effect of thioridazine against all the BC cells used in this study suggests that this drug could be used in association and/or as an alternative to Tam for the treatment of primary BC. Finally, new cellular pathways potentially druggable alone or in combination with ICI are being evaluated [13, 19, 20, 26, 27] as alternative strategies to treat metastatic BC. Because our ICI and thioridazine combination studies reveal a possible additive effect of these drugs in Tam-resistant MCF-7 cells, our data also suggest that this compound could be exploited as a co-treatment option for metastatic BC.

In conclusion, the present results confirm the anti-cancer activity of thioridazine [29] in BC cells modeling primary breast tumors (*e.g.*, MCF-7 and ZR-75-1 cells) and further extend this notion to BC cells mimicking Tam-resistant metastatic BC (*e.g.*, Tam-resistant MCF-7 cells and Y537S-MCF-7 cells). Moreover, our discoveries define, for the first time, the ability of thioridazine, a drug widely used in the clinic as an anti-psychotic, to reduce the ERα intracellular levels. The reduction in the ERα content in BC cells not only would most likely determine a reduced sensitivity to E2, which is fundamental for the progression of primary breast tumors but also would lead to the elimination of one of the main drivers (*i.e.*, ERα) of the uncontrolled proliferation of metastatic breast cancers.

Therefore, our study renews hopes for patients with BC by suggesting a re-purposing use of thioridazine for primary breast tumors as well as for metastatic Tam-resistant BC, including those diseases expressing the Y537S (and possibly others) ERα variants.

## Acknowledgments

This study was supported by grants from Ateneo Roma Tre to FA and from MIUR grant (Ricerca Corrente) to FB. The Grant of Excellence Departments, MIUR (ARTICOLO 1, COMMI 314 - 337 LEGGE 232/2016) to Department of Science, University Roma TRE is also gratefully acknowledged. The Authors thank also Prof. Simak Ali, University of London Imperial College, England for the gift of the Y537S-MCF-7 cells and Dr Carol Dutkowski, University of Cardiff, England for the gift of the 4OH-tamoxifen resistant MCF-7 cells (Tam-Res). We thank Dr Tommaso Colangelo and Orazio Palumbo, IRCCS Casa Sollievo della Sofferenza, Italy for the technical support for the affymetrix experiment. The Authors declare no conflict of interests.

